# Decomposing representational drift across wake and sleep

**DOI:** 10.64898/2026.03.31.715372

**Authors:** Julia J. Harris, Andreas Schaefer, Mihaly Kollo

## Abstract

Neural representations evolve over time, yet the relative contributions of online experience and offline states such as sleep remain unclear. Here, we recorded single-unit activity in the olfactory cortex of mice across cycles of awake odour exposure and sleep, and developed a low-rank decoder to track representational drift. We identified four orthogonal drift modes operating on distinct timescales, revealing that sleep and wake drive qualitatively different transformations, which indicates that offline reorganisation is not a simple continuation of online learning. Rather, sleep initiates an about-turn in the overall drift trajectory, which is uniquely characterised by a combination of decorrelation and rotation of odour representations. We also provide the first evidence for olfactory replay, occurring at ~2.5× temporal compression and associated with locally generated piriform cortex sharp waves. Together, these findings demonstrate that representational drift comprises state-dependent components, and reveal distinct contributions of wake and sleep to sensory representational change.

## Introduction

The brain’s representation of the world is not fixed but flexibly adapts to incorporate acquired information and skills (Bostock et al., 1991; Nudo et al., 1996; Poort et al., 2015; Recanzone et al., 1993; Schoups et al., 2001; Zohary et al., 1994), which are then consolidated into memory (Dudai et al., 2015; Kandel et al., 2014; McClelland et al., 1995; McGaugh, 2000; Scoville & Milner, 2000). Over a timespan of years, memories must be maintained by a neural code that is constantly challenged with new information (Frankland & Bontempi, 2005; Fusi et al., 2005; McClelland et al., 1995; Roxin & Fusi, 2013). The biological mechanisms underpinning both memory acquisition and long-term maintenance operate on much quicker timescales (e.g. synaptic plasticity and protein turnover on the order of seconds to minutes; Bliss & Collingridge, 1993; Frey & Morris, 1997; Kandel et al., 2014; Neves et al., 2008), yet must support stability in perception and behaviour (Acker et al., 2019; Susman et al., 2019).

As such, it has been surprising to discover over recent years that the neurons which encode even low-level sensory information change identity over time, a phenomenon called representational drift (Clopath et al., 2017; Deitch et al., 2021; Driscoll et al., 2017; Rule et al., 2019; Schoonover et al., 2021; Ziv et al., 2013). In the primary olfactory cortex, odour representations can be formed and stabilised quickly (Bolding et al., 2020), but then drift slowly over weeks to months (Schoonover et al., 2021). Whether such drift is a correlate of continued learning (Alevi et al., 2026; Driscoll et al., 2017) or a by-product of behavioural variability (Sadeh & Clopath, 2022) or stochastic synaptic fluctuations that must be compensated for (Micou & O’Leary, 2023; Rule & O’Leary, 2022) is not currently clear (Alevi et al., 2026).

Additionally, experimental measures of drift have not yet teased apart the relative contributions of wake and sleep. Sleep is increasingly recognised as a critical phase for memory consolidation, where offline reactivations of neural ensembles that were recently active during learning drive lasting improvements in behaviour (Bendor & Wilson, 2012; Born & Wilhelm, 2012; Dave & Margoliash, 2000; Diekelmann & Born, 2010; Ji & Wilson, 2007; Wilson & McNaughton, 1994). Sleep is also thought to mediate homeostatic regulation of neural energy use and coding efficiency (Tononi & Cirelli, 2014; Xu et al., 2024). Both of these functions would be underpinned by biological mechanisms operating on fast timescales (e.g. synaptic weight changes and firing rate renormalisation).

Thus, slow representational drift must necessarily arise from the accumulation of many of these shorter processes (Alevi et al., 2026), whose computational function (e.g. learning, consolidation, reorganisation) and physiological expression (e.g. adaptation, EI balance, spine turnover, firing and synaptic normalisation) may differ between awake and sleep states (Aton et al., 2009; de Vivo et al., 2017; Hengen et al., 2016; Sawada et al., 2024; Seibt et al., 2012; Suppermpool et al., 2024; Torrado Pacheco et al., 2021; Vyazovskiy et al., 2008; Xu et al., 2024). We decided to investigate whether representational drift over one behavioural unit of wake-sleep-wake could be decomposed to reveal distinct components operating over different timescales and mediating different changes.

We recorded single-unit activity in olfactory cortices across two periods of awake odour stimulation and an intervening period of sleep (Fig. S1). We built a low-rank odour decoder which identified four orthogonal drifts in odour representation operating on distinct timescales. Sleep initiated an about-turn in the compound drift trajectory, which was characterised by a unique combination of rotation and decorrelation of odour representations. Additionally, we found the first evidence of olfactory replay during sleep, occurring at ~2-3× temporal compression and associated with locally generated piriform cortex sharp waves. Together, these findings demonstrate that slow representational drift is composed of shorter state-dependent components, revealing distinct contributions of wake and sleep to sensory representational change.

**Supplementary Figure 1.**
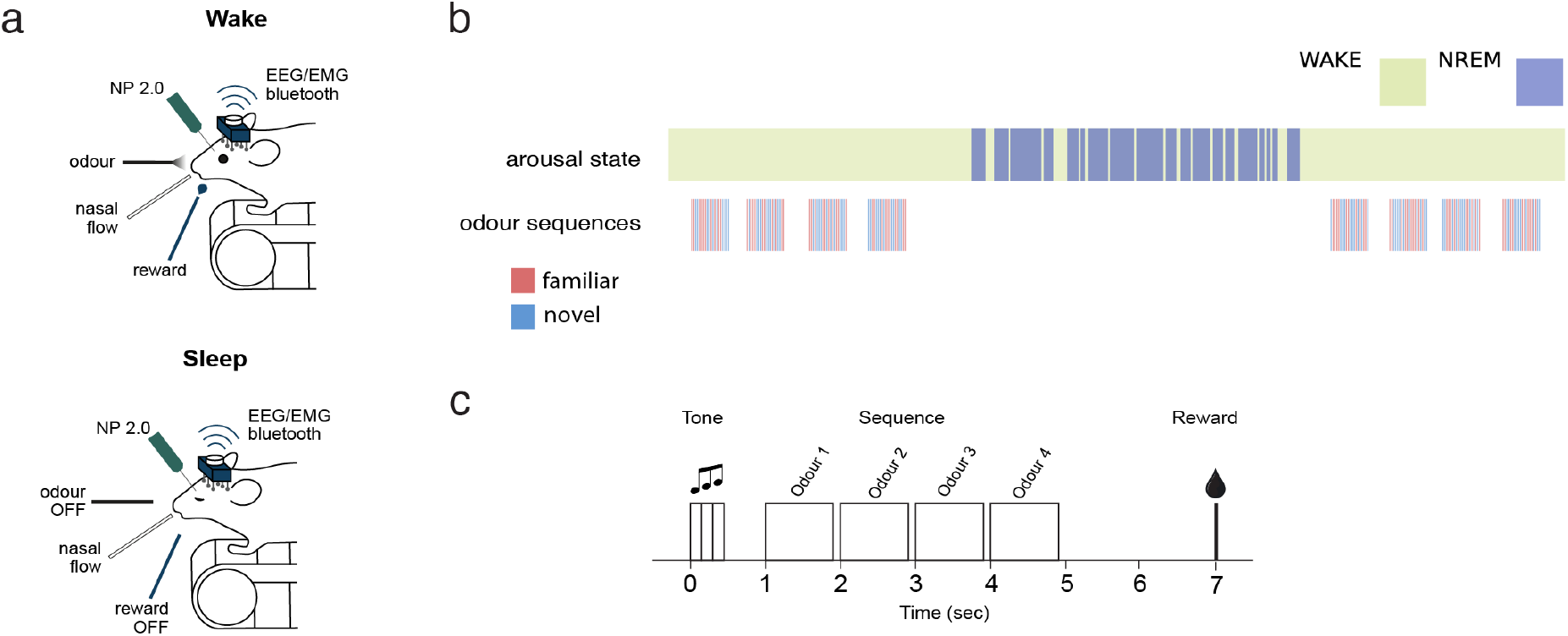
Experimental paradigm. (a) Head-fixed, simultaneous single-unit, EEG, EMG and respiration recording in the awake and sleeping state. Structure of the experimental session. 4 blocks of 40 odour sequences (familiar and novel) were played to the mouse followed by an approx. 1 hour long natural sleep followed by wake-up and a second group of odour blocks. (c) Stimulus trial paradigm: a cue sound was followed by a sequence of four unique odours (0.9s) every 1 s. Each odour was followed by a reward. Trials were separated by 30 s.

## Results

We performed single-unit recordings from neurons in the olfactory cortices of mice using NP2.0 multi-shank probes. EEG and EMG signals were simultaneously recorded to track vigilance state, and respiration was continuously monitored (Fig. S1a). During the first awake phase, mice were presented with novel and familiar four-element odour sequences (Fig. S1b, c). They were then allowed to fall asleep naturally, during which time no odours were presented. In the second awake phase, mice were again presented with the same odour sequences. Only units that were stable across the entire recording session were used for analysis (total of 636 units across 4 mice, localisation and curation described in Methods).

### Drift in olfactory representation can be decomposed into orthogonal modes operating at different timescales

Representational drift is known to occur in the piriform cortex on a timescale of weeks to months (Schoonover et al., 2021), but whether this drift is uniform across time is not known. Given that several of the candidate biological mechanisms for representational change (synaptic turnover, Hebbian plasticity, firing rate homeostasis) operate on much faster timescales (seconds to minutes), and that sleep is proposed to capatilize on some of these mechanisms to consolidate learning (Aton et al., 2009; Frank et al., 2001a; Seibt et al., 2012; Yang & Gan, 2012), we decided to look for processes contributing to representational change between the two awake periods (separated by an hour of rest).

To identify the different components of representational drift relevant for odour identification, we built a low-rank drift decoder W(τ) = W_0_ + ∑i αi(τ) · ΔWi, where each drift mode ΔWi is a low-rank matrix that tracks time-varying changes in the population readout while preserving stable stimulus identity (see Methods). The optimal decoder before sleep was broadly similar to the optimal decoder after sleep, showing that the neurons mostly maintained their tuning over this period (Fig. 1a). However, subtracting the final from the initial weight matrix revealed fine-grained changes distributed across both piriform and non-piriform neurons, indicating subtle reshaping of the neural code between awake phases (Fig. 1a, b). This difference could be decomposed into four largely orthogonal drift modes (*D1-D4*), each with a distinct sigmoidal time course that the model was free to learn independently (Fig. 1b, c). The four modes captured drift associated with initial odour exposure (*D1*), the wake-sleep transition and sleep itself (*D2 & D3*), as well as post-sleep exposure (*D4*).

**Figure 1.**
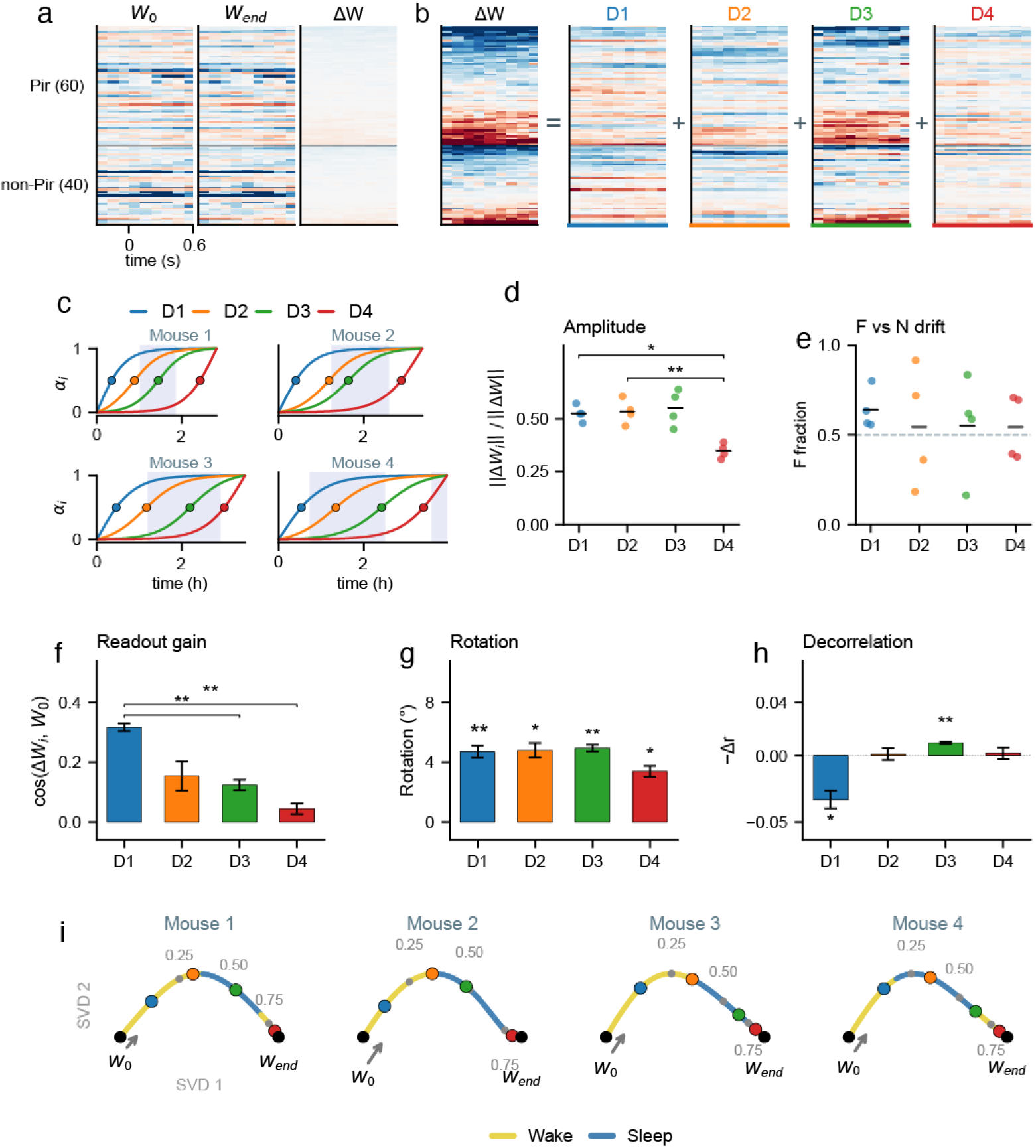
Odour representations undergo distinct transformations during wake and sleep. **(a)** Decoder weight matrices for a single odour (odour 1 of sequence F, mouse 2), showing similar baseline weights *W*_0_ and end-of-session weights *W* _*end*_. Their difference Δ*W* corresponds to a small change. Neuronal units sorted by mean Δ*W* per brain area. The colour scale (blue-white-red) is for negative to positive weights (identical colour scale used for all matrices in the panel). **(b)** The total change decomposes into four different drift modes *D1-4*. (Identical colour scale used for all matrices in the panel). **(c)** Normalised sigmoid activations for each mode in four mice. D1 (blue) activates during early wake, D2 (orange) and D3 (green) during the wake-sleep transition and sleep, and D4 (red) in post-sleep. Blue shading indicates the main NREM sleep block, odour sequences were played to the mice before and after sleep. Relative amplitude of each mode. Bars: mean ± SEM across 4 recordings. ^*^p < 0.05, Holm-corrected paired t-tests between mode pairs. **(e)** Fraction of each mode’s drift amplitude attributable to the familiar (F1) sequence odours. All modes affect both sequences approximately equally. **(f)** Readout gain: cosine similarity between each mode’s weight change and the baseline weights. D1 is significantly aligned with *W*_0_, indicating it scales the existing readout pattern. Stars: one-sample t-test vs zero, Bonferroni-corrected across 4 modes. **(g)** Rotation angle between *W*_0_ and *W*_0_ + Δ*W*_*i*_, in degrees. All modes produce significant rotation (^**^p < 0.01). **(h)** Odour decorrelation: change in mean pairwise Pearson correlation between odour weight vectors when each mode is applied (positive = increased distinctness). **(i)** Drift trajectory through weight space for each recording across the session. The four weight vectors Δ*W*(τ) are projected into 2D via SVD. The total drift direction is aligned horizontally. Coloured dots mark drift mode midpoints. Tick marks show τ values at 0.25 intervals. All mice show a similar path: D1 starts roughly along the baseline direction, then D2/D3 bend the trajectory during sleep, with D4 adding a minor late component.

We next asked what each drift mode learns about changes in the neural code. *D1*, active during initial odour exposure was largely aligned with the baseline readout weights (cosine similarity 0.32, Fig. 1f), increasing the readout weights in the direction of the existing code and increasing the pairwise correlations between odour responses (Fig.1h; p=0.02). Together, these indicate that odour representations became less distinct during the first presentations, consistent with adaptation reducing neural contrast, while the decoder compensated by amplifying its weights to maintain discrimination. In contrast, *D2-D4* were near-orthogonal to the baseline (Fig. 1f), indicating that later drift modes introduced genuinely new directions in weight space, rather than scaling existing tuning. All four modes produced comparable rotation of the readout vector (4-5° per mode, Fig. 1g), showing that representational drift, in the sense of modifying which neurons are most informative for stimulus identity, occurs across all phases. The modes diverged most sharply in their effect on inter-odour similarity (Fig. 1h). *D1* significantly increased pairwise correlations between odour representations, consistent with the redundancy expected from adaptation. The sleep-associated mode *D3*, in contrast, drove significant decorrelation (p=0.003), reducing the similarity between population responses to different odours. This combination of rotation and decorrelation during sleep aligns with normative predictions from efficient coding theory (Barlow, 1961; Chalk et al., 2018; Clifford et al., 2001; Simoncelli & Olshausen, 2001), suggesting that sleep consolidation actively optimises the population code for discriminability, while the rotation component reflects a continuous reorganisation of neural tuning (Alevi et al., 2026; Driscoll et al., 2017), which might reflect learning in non-odour-identity-related dimensions.

### Sleep drives drift orthogonal to wake adaptation

By fitting the drift model, we obtained a continuous, high-dimensional trajectory of the population code from the start to the end of each recording. We visualized these trajectories by projecting them into two dimensions (accounting for 65–74% of the total variance across mice), which revealed a remarkably consistent geometry (Fig 1i): wake-associated drift proceeded along the dominant axis, while sleep drove the representation orthogonally. During sleep, modes D2 and D3 whose activation timecourses partially overlapped (Fig. 1c) jointly drove the representation along an arc orthogonal to the wake drift axis, producing the largest displacement from the initial code. Strikingly, after sleep, the trajectory nearly stalled: post-sleep wake contributed little further drift (D4), indicating that the representational changes induced by sleep and initial adaptation together account for nearly all of the session’s total drift.

### There is spontaneous replay of odour sequences during sleep

The above results reveal drift operations that change the representation of olfactory stimuli over sleep. Given that both wake (Liu et al., 2025; Nguyen et al., 2024) and sleep (Bollmann et al., 2025) reactivation events have been associated with representational change, we hypothesised that olfactory replay should be evident across the same time period. Running the decoder across the entire length of the recording (not just during stimulus presentation), revealed odour predictions from spontaneous activity during sleep, which followed the order of awake-presented sequences (Fig 2a). To assess the significance of these putative replay events, we used temporal delayed linear modeling (TDLM, Liu et al., 2021). First, we built an empirical transition matrix between odours decoded from spontaneous activity during sleep and wake (Fig. 2b). In comparison to a circular-shifted null transition matrix, sequence-like transitions were significant for both novel and familiar sequences, across all recordings (Fig. 2c). Repeating this process at different compression factors revealed that sequenceness was maximal at a compression factor of 2-3x (Fig. 2d). Both partial (three-element) and complete (four-element) replay events were observed (Fig. 2e).

**Figure 2.**
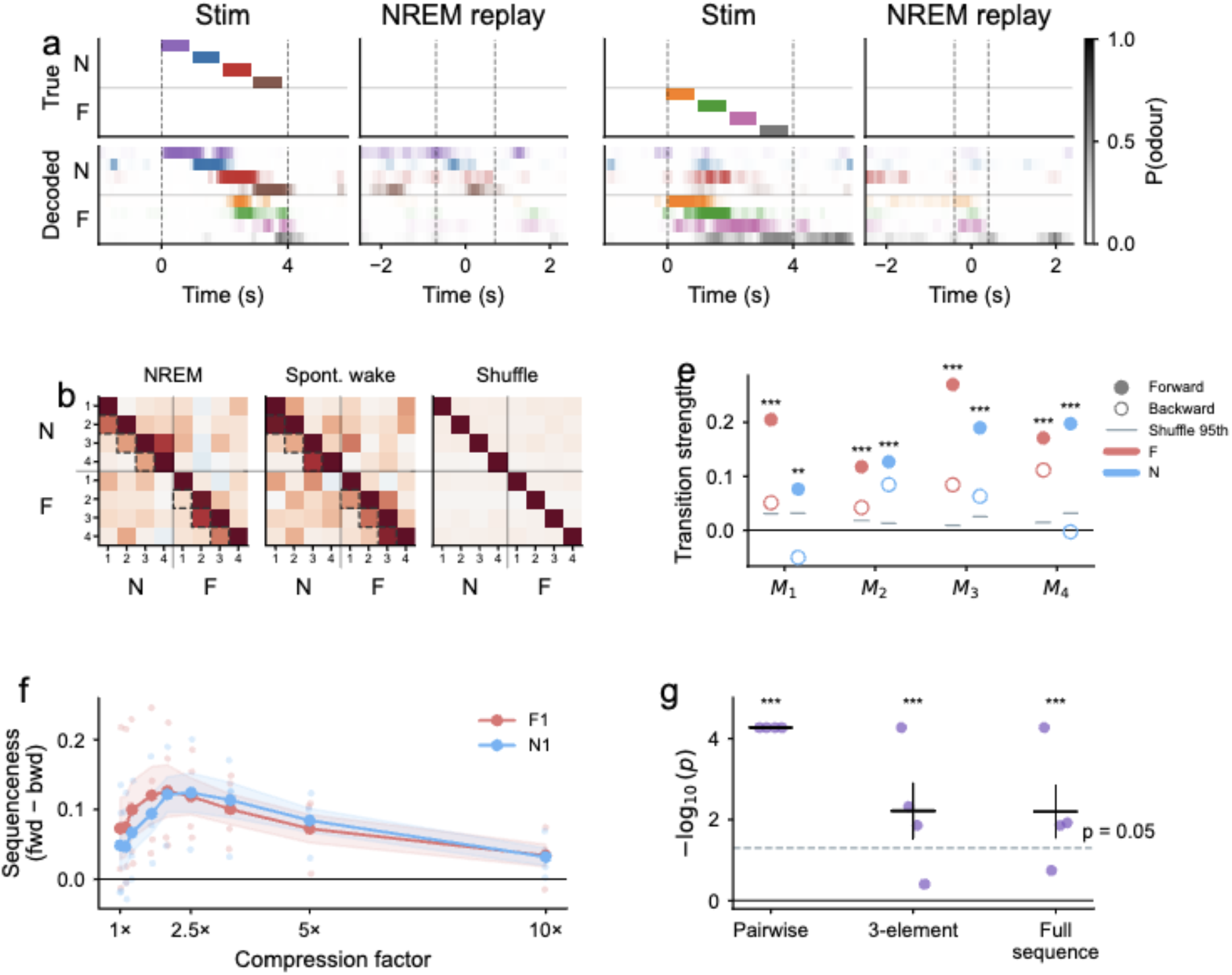
Temporally compressed odour replay during NREM sleep. **(a)** Example stimulus presentations (left) and NREM replay events (right) for the novel (N, left pair) and familiar (F, right pair) odour sequences. Top rows: true odour identity; bottom rows: decoded probability P(odour) from the drift decoder, displayed as colour-coded RGBA heatmaps (2-bin boxcar smoothed). Each row corresponds to one odour in the sequence; rows are grouped by sequence (N_1_-N_4_, F_1_-F_4_). Dashed lines mark stimulus onset/offset (stim panels) or first/last detected replay peak (replay panels). During wake stimulation, the decoder produces sequential activations matching the presented odour order. During NREM, spontaneous reactivations recapitulate the learned sequence at compressed timescales. **(b)** Transition matrices showing the regression weight from each odour’s decoded probability at time *t* to every other odour at time *t* + Δ*t* (ridge regression), computed at 2.5× temporal compression (Δ*t*=400ms). Dashed boxes highlight forward transitions along the trained sequence order. NREM shows stronger within-sequence forward structure than wake or circular-shifted channel shuffle (mean of 50 shuffles). **(c)** Forward (filled circles) and backward (open circles) transition strengths per mouse at 2.5× compression, for F1 (red) and N1 (blue) sequences separately. Horizontal grey lines show the 95th percentile of the circular-shift null distribution (500 shuffles). Forward transitions consistently exceed backward and shuffle in all mice. **(d)** Sequenceness (forward − backward transition strength) across temporal compression factors (1×-10×) for both sequences (F red, N blue). Lines: individual mice; shading: SEM. Peak sequenceness occurs at 2-3× compression for both sequences, corresponding to replay timescales of 300-500 ms per odour (vs 1 s during wake presentation). **(e)** Statistical significance of sequential replay structure at 2.5× compression, shown for three levels of sequence analysis: pairwise transitions (adjacent odour pairs), 3-element subsequences, and full 4-element sequences. Each dot is one mouse × sequence combination.

### Olfactory replay is associated with piriform cortex sharp-waves

In the hippocampus, replay events are closely associated with sharp-wave ripples (SWRs; Nádasdy et al., 1999). In the piriform cortex, similar population-level events have been described as sharp waves (SPWs), reflecting the absence of accompanying ripple oscillations (Katori et al., 2018; Manabe et al., 2011; Narikiyo et al., 2014). These events are not temporally coupled to hippocampal SWRs, but instead arise locally within the piriform cortex and propagate to other olfactory regions, raising the possibility that the piriform cortex independently coordinates olfactory consolidation during sleep. Having identified olfactory replay events for the first time (Fig. 2), we were in a position to test whether they are associated with piriform SPWs, as spatial replay events are with hippocampal SWRs.

First, we identified SPWs, which were defined as large negative deflections in the 5-20 Hz frequency band of the LFP (Fig. 3a). We found that these events were most common in the piriform cortex and their occurrence increased during sleep (Fig. 3b). In contrast to previous reports (Katori et al., 2018; Manabe et al., 2011; Narikiyo et al., 2014), however, we found that power in both the ripple band (160-190 Hz), and gamma band (80-140 Hz; generally elevated in piriform cortex) increased immediately after sharp-wave occurrence (Fig. 3c). The increase is subtle compared to that seen after hippocampal sharp-waves (Buzsáki et al., 1992), so we think that the term SPW (rather than SWR) is still appropriate to highlight the qualitative difference in population events, although we want to highlight that piriform cortex SPWs do comprise some high-frequency activity. Finally, we found that replay rate was modulated by proximity to the peak of SPWs (Fig. 3d), with an increased occurrence of replays within the 500 ms following SPWs (Fig. 3e), suggesting that, as in the hippocampus, these LFP events are associated with replayed content.

**Figure 3.**
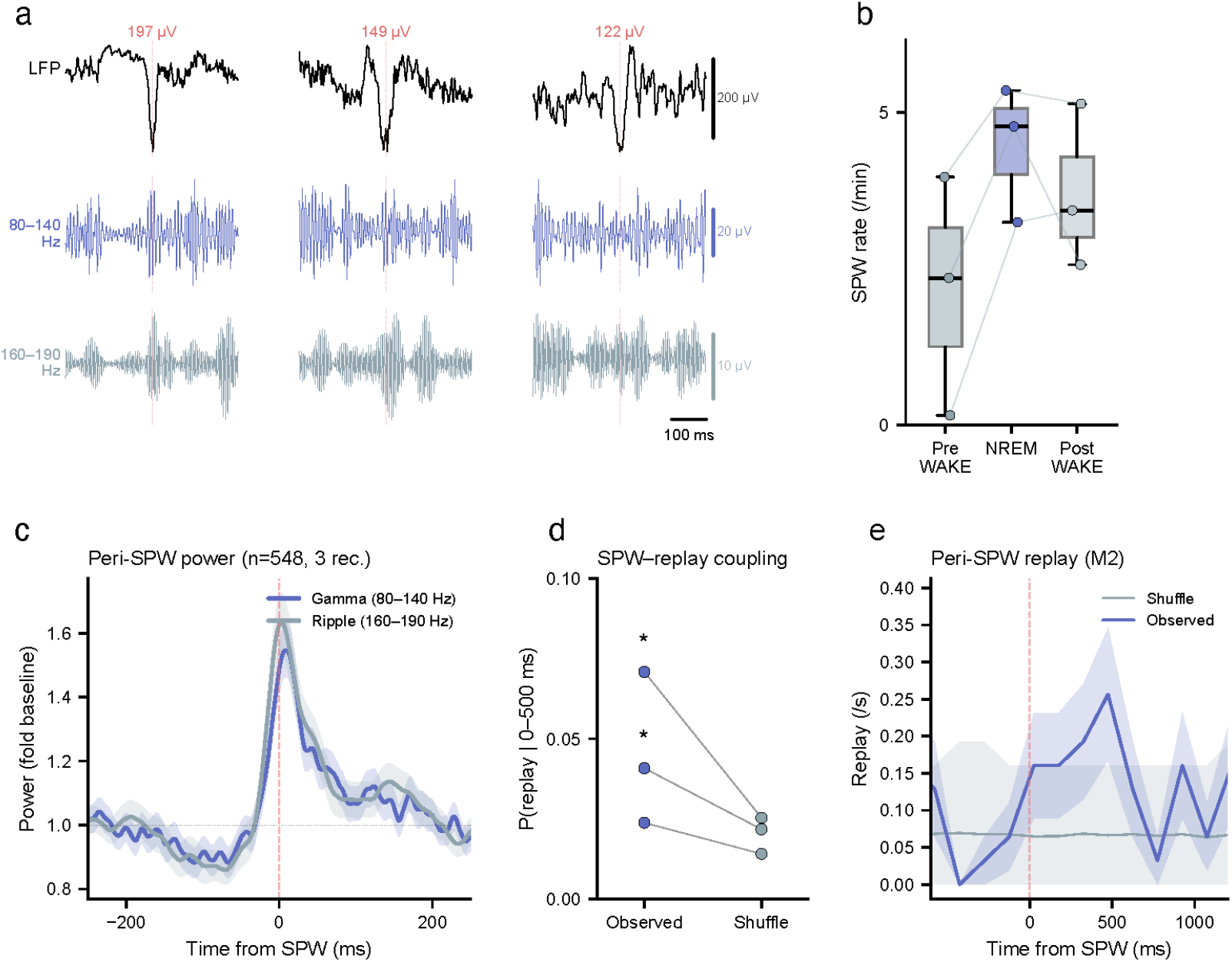
Piriform sharp waves during NREM sleep are associated with odour replay. **(a)** Example piriform sharp waves (SPWs) detected during NREM sleep. Top: raw LFP from a piriform channel showing the sharp wave deflection (negative polarity, peak amplitude annotated in µV). Middle: 80-140 Hz filtered signal showing co-occurring fast gamma activity. Bottom: 160-190 Hz filtered signal showing ripple-band oscillations. **(b)** SPW rate across behavioural states. Box plots show the distribution across recordings (n = 3) for pre-sleep wake, NREM, and post-sleep wake. Paired lines connect the same recording. SPW rate is elevated during NREM relative to wake periods. **(c)** Peri-SPW triggered power in gamma (80-140 Hz, blue) and ripple (160-190 Hz, grey) frequency bands. Both gamma and ripple power peak sharply at the time of the SPW, with gamma showing a ~60% increase and ripple ~15% above baseline. **(d)** SPW-replay coupling. The probability of observing a replay event (decoder P(odour) > 0.05) within 0-500 ms following a sharp wave, compared between observed data and circular-shift null (500 shuffles). Each dot is one recording. Two of three mice show significant SPW-replay co-occurrence, comparing observed coupling rate against the null distribution. (M1 p = 0.033, M2 p < 0.001). **(e)** Peri-SPW replay PSTH for the recording with the strongest coupling (M2). Blue: observed replay rate in 150 ms bins around each sharp wave. Grey: shuffle expected rate with 95% CI envelope. The observed replay rate peaks in the 0-500 ms window following the SPW, exceeding shuffle expectations.

## Discussion

We demonstrate that online and offline periods make independent, qualitatively distinct contributions to representational change in the neural code. Rapid drift was triggered by initial odour exposure, which followed an adaptation-like signature. Drift continued along the same broad trajectory until the animal was in full sleep. Across mice, sleep governed an about-turn in the compound drift trajectory. Sleep is when representations by olfactory cortical neurons were maximally rotated (odours decoded from a shifted neural ensemble) and decorrelated (reduced overlap between neurons responding to different odours). Subsequent wake and repeated odour exposure saw almost no further drift. These findings provide the first decomposition of compound representational drift into distinct components operating over different timescales. Critically, sleep-mediated changes did not simply continue the changes occurring during wakefulness, suggesting that wake and sleep replay - both observed here in the olfactory cortex for the first time - contribute differently to neural reorganisation.

### State-dependent components of representational drift

Neural representations of even low-level sensory stimuli drift slowly over time (Clopath et al., 2017; Deitch et al., 2021; Driscoll et al., 2017; Rule et al., 2019; Schoonover et al., 2021; Ziv et al., 2013); in the piriform cortex, the representation of odours change on the order of weeks to months (Schoonover et al., 2021). But such gradual drift must be the sum of biological processes that operate on much faster timescales (for example, spine turnover, Hebbian plasticity, synaptic normalisation or firing rate homeostasis, all operating on the order of seconds to minutes). These processes are unlikely to act uniformly across time, with differing dynamics particularly likely across cycles of waking and sleeping (Aton et al., 2009; de Vivo et al., 2017; Frank et al., 2001b; Hengen et al., 2016; Sawada et al., 2024; Seibt et al., 2012; Suppermpool et al., 2024; Torrado Pacheco et al., 2021; Vyazovskiy et al., 2008; Xu et al., 2024). However, teasing out different components and timescales of representational drift from experimental data has not been possible to date.

Here, we developed a drift decoder capable of decomposing representational change into distinct drift modes. Applying this to single-unit activity recorded from olfactory cortices across two periods of awake odour exposure with an intervening period of sleep revealed four orthogonal drift modes with different timescales. Substantial and unique changes occurred over sleep: odour representations were maximally rotated and decorrelated by this component. Additional drift modes described changes during initial odour exposure (larger weights were required to decode odours, and contrast was decreased); sleep onset (rotation and weight gain); and post-sleep odour experience (rotation with decorrelation). Across mice, the relative timing and trajectories of these drift modes were remarkably similar, and full sleep was always associated with an about-turn in the compound drift trajectory.

That we were able to observe any representational change over the course of minutes to hours may seem surprising given that the overall rate of retuning in the piriform cortex takes weeks to months (Schoonover et al., 2021) It is important to note that odour identity decoding did remain stable across our recordings (Fig. S2a), with only small changes to weights between the initial and final wake phases (see Fig.1a). This is exactly the point: drift that takes months to emerge at the population level must result from the accumulation of subtle drifts at short timescales. Our data and novel drift decoder have enabled the first detailed investigation disentangling these short-term changes at the network level.

Crucially, by identifying the exact transformations that these distinct drift modes govern, our data also highlight that sleep-mediated changes are not a simple continuation of wake changes; rather, sleep prioritises a unique set of transformations. Do these transformations correspond to existing proposed functions of sleep?

### Relationship between sleep drift and theories of sleep’s function

Sleep is widely considered to support memory consolidation as demonstrated by improving behavioural performance (Born & Wilhelm, 2012; Diekelmann & Born, 2010), and is also proposed to serve a homeostatic role, renormalising synaptic strengths (de Vivo et al., 2017; Sawada et al., 2024; Suppermpool et al., 2024; Tononi & Cirelli, 2014; Vyazovskiy et al., 2008), stabilising firing rates (Torrado Pacheco et al., 2021), and restoring network-level properties such as criticality (Xu et al., 2024). Do the representational changes described by the main sleep drift mode observed here - rotation and decorrelation - correspond to any of these ideas?

Interestingly, rotation is representational drift’s canonical transformation; where a different set of neurons become the most informative about the same stimulus. An open question is whether rotational drift is a form of memory consolidation (for example, geometry changes to make space for the storage and integration of new memories), or whether it arises as a natural consequence of ongoing structural remodelling (such as synaptic turnover), which may be enhanced during sleep. The passive nature of odour exposure in the present experiment limits our ability to differentiate these possibilities, but a clear next step will be to employ a task with increased complexity and learning demands. For example, requiring the animals to alternately categorise odours as similar or different (Bostock et al., 1991) would enable quantitative predictions about how representations should reorganise, which the drift decoder could track across wake and sleep phases. However, even without deciphering the computational reason for these rotations, our findings deliver an important message: the change in drift trajectory between initial odour exposure and sleep, along with the observation that awake experience tends to slow rotational drift (Schoonover et al., 2021), contribute to a picture that offline reorganisation may be the primary driver of slowly evolving representational drift

While some degree of rotation is present in all drift modes identified, decorrelation is unique to sleep. Sleep reduces the overlap between neurons representing different odours, increasing discriminability in the neural code. Comparable effects have been observed in visual cortex as a function of awake reactivation rate (stimulus response orthogonalisation; Nguyen et al., 2024), and in prefrontal cortex after NREM sleep (cortical circuit desynchrony; Kharas et al., 2024). Decorrelation suggests a consolidation role for sleep that also serves homeostatic efficiency. Piriform cortex exhibits highly selective odour responses (Bolding & Franks, 2017, 2017; Davison & Ehlers, 2011; Kehl et al., 2024; Poo & Isaacson, 2009; Stettler & Axel, 2009), and our data indicate that while initial odour exposure increases correlation between odour representations and decoder weights required for classification, sleep strategically reorganises the system to ensure distinct neural ensembles represent different odours. This coding arrangement maximises storage capacity (Marr, 1971; Olshausen & Field, 2004; P Foldiak & M Young, 1995) while minimising interference (McClelland et al., 1995) and energy costs (Lennie, 2003; Levy & Baxter, 1996).

Thus, sleep mediates large changes in neural representation, which are qualitatively different to awake transformations, and fit with both consolidation and homeostatic theories of sleep.

### Local replay, sharp-waves and drift

We report the first evidence of olfactory replay, with a predominant playback speed of ~2-3x compression. This finding adds to a growing body of work demonstrating replay across multiple modalities beyond the classic spatial hippocampal replay (Dave & Margoliash, 2000; Eagleman & Dragoi, 2012; Ji & Wilson, 2007; Liu et al., 2019; Peyrache et al., 2009; Thompson et al., 2024). The association of olfactory replay events with piriform cortex-generated sharp-waves, which are known to be independent of hippocampal sharp-wave ripples (Katori et al., 2018; Manabe et al., 2011; Narikiyo et al., 2014), also contribute to an emerging picture that replay can be locally generated, without hippocampal coordination (Thompson et al., 2024).

Presumably, local replays contribute to local network reorganisation. An obvious question is whether the olfactory replay events that we observe are responsible for driving the measured sleep drift. Recent work in the visual cortex demonstrated that awake reactivations track the orthogonalisation of stimulus representations, and predict future stimulus responses (Nguyen et al., 2024). In the hippocampus, reactivated assembly patterns during sleep gradually transform to resemble those seen in future waking (Bollmann et al., 2025). In the olfactory bulb, electrical stimulation during sleep to mimic reactivation augmented fear memory (Barnes & Wilson, 2014). It thus seems likely that olfactory replay drives at least a portion of sleep drift, but other features such as sleep depth or duration could also be responsible, and important next steps are to tease out the respective contributions of these phenomena. The observed coupling between replay and sharp-waves make causal intervention tractable: closed-loop SPW manipulation (similar to that done in the hippocampus, (Fernández-Ruiz et al., 2019; Girardeau et al., 2017)) will allow for direct assessment of their impact on each identified drift mode.

### Summary

Together, our findings reveal that representational drift is not a uniform process but comprises multiple, state-dependent components that unfold over distinct timescales, with different contributions to overall changes in representation. In particular, sleep is a period of substantial network reorganisation characterised by representational rotation and decorrelation. These changes are distinct from the transformations that happen during awake experience, consistent with a role for sleep in actively rebalancing and restructuring sensory representations rather than simply reinforcing prior changes. The discovery of local olfactory replay suggests that such circuit transformations can be mediated locally, and we hypothesise that similar processes are occurring across the brain during sleep, without the requirement of a central coordinator. These results provide a framework for linking rapid, state-dependent plasticity to long-term representational change, and establish a foundation for future causal tests of how replay and sleep shape the evolution of sensory codes over time.

## Materials and Methods

### ANIMALS

#### Animal handling

All procedures were carried out in accordance with the Animals (Scientific Procedure) Act of 1986. Animal research was approved by the United Kingdom Home Office and the Animal Welfare and Ethical Review Panel of the Francis Crick Institute. Male mice (C57BL6/J) were kept on a standard 12-h/12-h light/dark cycle (lights on at 10 am) with standard chow and water available ad libitum throughout the study. Each mouse underwent two surgeries: the first to chronically implant EEG/EMG and headplate, and the second to perform a craniotomy for acute neuropixels recordings. Mice were housed together with their littermates up until the day before the second surgery (so each mouse was singly housed for its experimental week). Experiments were initiated at the transition from dark to light, to align with the natural phase of high sleep pressure.

#### Surgical procedures

All surgeries were performed with mice in a stereotaxic frame under isoflurane anaesthesia (~1.5%), with meloxicam (2 mg/kg of body weight) and buprenorphine (0.1 mg/kg of body weight) delivered s.c. for analgesia. The first surgery (EEG/EMG and headplate) was performed when mice were at least 16 weeks old and weighed 26-30g. After shaving the head, the skin was cleaned with chlorhexidine (0.5%). Next, scissors were used to create an incision in the skin from above the nuchal muscles to above the olfactory bulb. The skull was cleaned and skin was held open with small Serrefine Bulldog Clamps (Fine Science Tools). Four miniature EEG screw electrodes were then implanted (from bregma: AP +1.5 & ML −1.5 (common reference); AP −1.5 & ML −1.5 (EEG1); AP −1.5 & ML +1.5 (EEG2); AP −3 & ML −2 (ground)), and two EMG electrodes were inserted into the nuchal musculature. These electrodes were each connected, via an enamel-coated copper wire, to an individual gold pin of an EEG/EMG headstage (Harris et al., 2022), which was mounted onto a custom steel headplate. The headstage/headplate combo was positioned directly above lambda using a custom stereotaxic holder, and then lowered and tissue-glued to the skull. Coordinates for future craniotomy (second surgery) were marked above the piriform cortex (from bregma: AP +1.3 & ML +3) and dental cement (C&B Superbond) was carefully applied to form a circular well evenly centred around the piriform cortex coordinate, with diameter ~4 mm. Dental cement was then used to cover all EEG screws and wires, and secure the headstage/headplate in place. Skin was drawn over the dental cement well and sutured around the headplate, and animals were returned to their home cage, with their littermates. After recovery (≥7 days) plus sufficient habituation to head-fixed sleep (≥10 days, see below), the second surgery was performed.

The day before the second surgery (craniotomy for acute neuropixels recordings), mice were singly housed. During this surgery, scissors were used to open the skin above the pre-formed dental cement well. A circular craniotomy (~2 mm diameter) was made in the centre of the well (i.e. at the previously measured piriform cortex coordinates), and a durotomy performed with a blunt needle. The craniotomy was covered with low melting point agarose, and a layer of Kwik-Cast was added after the agarose had set (with a thin film of saline between the brain and the agarose, and the agarose and Kwik-Cast). The skin was sealed to the outside of the dental cement well using tissue adhesive, and the animal was returned to its home cage.

#### Habituation to headfixed sleeping

After recovery from the first surgery, mice were trained to sleep while headfixed. Mice were habituated to headfixation for a minimum of ten days, progressing through the following steps: gentle handling; brief bouts of headfixation (from 5 sec to 50 minutes) in their holding room (using a portable stage) with hand-delivered soy milk rewards; up to one hour of headfixation in the recording rig with pump-delivered soy milk rewards, and then the addition of continuous clean air delivery to the nose. The final phase of habituation introduced cues to signal a rest period, which were employed in every experiment thereafter: an ambient light was turned on and a black curtain around the rig was drawn closed. This phase was conducted while recording EEG/EMG to confirm that the mouse slept during the rest period. The headfixation stage consisted of a high-friction, flat treadmill, meaning that running took some effort and staying still felt stable. The head was positioned relatively low with respect to the body so that neck and limbs were not elongated, and a custom covering was positioned over the body of the mouse, to provide warmth and comfort. Headfixation in the rig was always timed to commence within the first hour of the mouse’s light phase, to align with high sleep pressure. Once a mouse was able to sleep while headfixed, it entered the pipeline for its experimental week.

## EXPERIMENTAL PROCEDURES

### Experimental pipeline

#### Craniotomy day (Day 0)

On the morning of Day 0, the mouse was headfixed in the recording rig (air table, Faraday cage), and introduced to a sequence of four odours, which would thereafter be the “familiar sequence” (F1) for that mouse (see Odour details below). The mouse was then allowed to rest while headfixed for approximately 60 minutes (sleep monitored using EEG/EMG), and then returned to its home cage for 1-2 hours. On the afternoon of Day 0, the mouse underwent surgery two (piriform cortex craniotomy), in preparation for acute neuropixels recording the next day (Day 1).

#### Experimental day (Day 1)

The mouse was headfixed and the AP/ML axes were adjusted using a bullseye bubble spirit level custom-mounted to an EEG insert, which was subsequently replaced with the wireless Bluetooth EEG/EMG amplifier (Pinnacle, see below). The layer of Kwik-Cast was gently removed, and the multishank neuropixels probe (NP 2.0) was coated in a lipophilic dye (DiI, DiO or DiD, different for each day) and then inserted through the layer of agarose. To follow along the anterior-posterior axis of the piriform cortex, the probe was angled at 20° from the AP axis, and to accommodate the slight curvature of the piriform cortex at the base of the brain, the probe was angled at 17° from the DV axis (Fig. S3a). Coordinates were planned using Pinpoint (Virtual Brain Lab; https://virtualbrainlab.org/pinpoint/). We lowered the probe (movement along the micromanipulator’s axis of travel, Luigs & Neumann), to an approximate depth of 3.5 mm, estimated from the surface of the brain. Because the surface point was hard to be precise about through the agarose, we thereafter relied upon electrophysiological features of the piriform cortex as we lowered the probe to its final depth (M1 −3.82; M2 −3.45; M3 −3.90; M4 −3.92). After a 20-30 min stabilisation period, mice entered the first odour stimulation phase of the experiment, followed by the rest phase, and then the second odour stimulation phase (details below). Each phase lasted approximately 1 hour, meaning that the total experimental time was 3-4 hours. The probe was then slowly removed, and cleaned with warm Tergazyme (Alconox) followed by distilled water. At the end of the experiment, the mouse was transcardially perfusion-fixed and the brain extracted for histology (details below).

### Awake odour stimulation

Odour stimuli were sequences of four odours, presented for 900 ms each with 100 ms between odours (Fig S1c). Each odour sequence was preceded by a unique tritone played from speakers placed on either side of the mouse (Logitech), and followed by a soy milk reward which was not contingent on stimulus or behaviour.

Stimuli were presented in 20-trial blocks, with sequences presented every 30 s (totalling 10 min). For blocks with more than one sequence type, equal numbers of each type were presented within each block, delivered in a randomised order. Typically, four blocks were presented before and after sleep. Blocks were separated by 5 minute rest periods, during which the animal typically displayed quiet wakefulness, and rarely slept.

Sequence types were changed systematically across the three experimental days to vary different aspects of novelty (see Table 1). Odours for each sequence were counterbalanced across mice: half of the mice had the opposite odour sequences pictured in (i.e. familiar and novel sequences and their derivatives were flipped).

**Table 1.** Odour stimulus identities. IDs are arbitrary codes based on Canberra distances in chemical space (e.g. C3f = Cluster 3, far).

| Odour ID | Chemical |  |
| --- | --- | --- |
| C3f | Methyl benzoate |  |
| C4f | Cineole |  |
| C4cA | Citronellol |  |
| C4cB | Geraniol |  |
| O | 2-Hydroxyacetophenone |  |
| MeA | Alpha terpenine | Ethyl caproate |
| MwA | Methyl acetophenone | Cyclohexanol |
| MwB | Acetophenone | Guaiacol |

### Odours

Eight different odour stimuli were used: five were monomolecular odours and three were 1:1 mixtures of two monomolecular odours (Table 1). Odour choices were made as follows: (1) Mordred molecular descriptors were used to map 82 odours onto a projection of chemical space, and selected odours must have large Canberra distances from each other; (2) the range of possible ppm values that could be achieved by air dilution via high speed valve cycling was calculated based on the vapor pressure of each odour, and selected odours must be presentable within 5-fold of the target ppm of 10 (range: 7-51 ppm); (3) odours must not be known to be innately attractive, aversive or appetitive to mice.

### Odour delivery

Odours were delivered via a custom-built, all-air-dilution 16-channel olfactometer (based on Erskine et al., 2019). Odour channels contained pure chemicals in individual glass bottles, while four blank channels housed empty bottles. Each bottle was flanked by two valves: an upstream “back valve” (NO or NC 3-way solenoid valves, OEM Automatic) to prevent diffusion-based contamination of clean air input, and a downstream “front valve” (VHS Series 2-way solenoid valves, Lee Company) to control release into one of four parallel manifolds, which converged to a single output path to the mouse. Between odour presentations, a continuous carrier stream flowed through the blank channel on each manifold (0.2 ml/min per channel; 0.8 ml/min total). During odour delivery, one blank channel was closed and one odour channel opened, maintaining constant total flow.

Air dilutions were calculated for each odour based on inherent vapor pressure and target ppm (see above) and achieved by high-speed valve cycling at a custom duty cycle for each odour, which was flow-compensated by equal and opposite cycling of the blank valve on the same manifold. Due to low-pass filtering through the tubing (odour-resistent MINSTAC clear teflon, Lee Company), the odour stimulus measured at the position of the mouse’s nose using a photoionisation detector (mini-PID, Aurora Scientific) were smooth pulses (i.e. did not reflect valve cycling).

The olfactometer was housed in a sound-insulated box. It was supplied by compressed air, which was filtered (SMC AMF250C series 5 μm, 0.1 MPa to 1 MPa Pneumatic Filter, RS Components) and mass-flow controlled (SMC G 1/8 Pneumatic Regulator −0.05 MPa to 0.85 MPa, RS Components). Flow was additionally controlled at every individual channel via a regulator before each back valve (FR2000 Series Variable Area Flow Meter for Gas, 0.1 L/min Min, 1 L/min Max, Key Instruments). The olfactometer’s output was delivered to the head-fixed mouse via a specialised three-way nosepiece made of odour-resistant Teflon (design generously provided by Bolding & Franks, 2018), which allowed air to flow across the nose and respiration to be monitored simultaneously via a flow meter (details below). Exhaust was vented to the facility’s central vacuum extraction.

### Rest/sleep period

After the first phase of awake odour stimulation, the habituated cues to signal a rest period were deployed (black curtain and ambient light, described above) and odour stimulation ceased. The rest period was typically 60-90 minutes, guided by continuous monitoring of the EEG/EMG and neuropixels data. We allowed for natural variability in sleep amount during this time. At the end of the rest period, we opened the curtain, turned off the ambient light, and resumed odour stimulation.

## DATA ACQUISITION AND SYNCHRONISATION

### EEG/EMG

For both home cage and headfixed vigilance state monitoring, EEG and EMG signals were recorded using Pinnacle’s 3-channel wireless system (8200-K9-SL; Pinnacle Technology Inc). Signals were high pass filtered by the preamplifier (0.5 Hz for EEG and 10 Hz for EMG) and then recorded using Pinnacle’s Sirenia Acquisition Software, sampled at 1024 Hz.

### Neuropixels

For acute, headfixed Neuropixels recording, signals were acquired at 30 kHz using the PXIe base station (Imec / National Instruments) and OpenEphys acquisition software. LFP signals were band-pass filtered at 0.1-500 Hz, and spike signals at 300-6000 Hz. As the piriform cortex is positioned ventrally, we recorded from the bottom 48 channel rows on each shank. We used an external ground configuration: the ground and reference pads were connected and then led by a single wire to the implanted ground screw (same screw as for the EEG ground).

### Olfactometer

The olfactometer was controlled via custom software (vscode / Python) which interfaced with National Instruments hardware (cDAQ-9178) and custom-designed PCBs (linked with NI-9923) to command individual back-valves (NI-9477) and front-valves (NI-9403) with concerted timing.

### Respiration and licking

During headfixation, respiration was continually measured using a flow meter (Honeywell Amplified Airflow Sensor, RS Components), and licking was detected by a piezo-based sensor. Both signals were acquired using Spike2, via a CED Micro 1401 acquisition unit (Cambridge Electronic Design Ltd), with a sampling rate of 2 kHz.

### Synchronisation

The CED Micro 1401 was used as the central clock, with regular synchronising signals from the EEG hardware, Neuropixels hardware and the olfactometer hardware all combined in a single Spike2 master recording for each experiment, with the aid of a custom-written Bonsai workflow (https://bonsai-rx.org/).

## DATA ANALYSIS

### Vigilance state scoring

Sleep states – NREM, REM and wake – were automatically classified and then manually verified from EEG and EMG data as in (Harris et al., 2022). Wakefulness was defined as de-synchronised, low amplitude EEG and tonic EMG with bursts of movement. NREM sleep was defined as synchronized, high amplitude EEG in the delta frequency range (1-4 Hz) and reduced EMG activity relative to wakefulness. REM sleep was defined by high theta:delta ratio with an absence of muscle tone, but was rarely seen in the headfixed sessions (because they occupied the first hour of the day’s sleep, when REM sleep is not prevalent). To be sure that mice could enter true sleep while headfixed, we performed these tests in an initial group of four mice who also had neuropixels recordings but did not receive the same odour stimulus sets, as well as the four mice used for the full experiment here (Fig. S4).

### Spike sorting and curation

Neuropixels 2.0 recordings were spike-sorted using Kilosort 4 with default parameters. Raw data were preprocessed with common average referencing per shank and bandpass filtered (300–6000 Hz) prior to sorting. Spike sorting was performed with Kilosort 4 (version 4.0) using conservative default parameters: spike detection thresholds of 9 (universal templates) and 8 (learned templates), non-rigid drift correction (nblocks = 1), cross-correlogram merge threshold of 0.25, and batch size of 60,000 samples. particularly during periods of drift in spike amplitude or waveform shape, as occurs across sleep-wake transitions, rather than risk merging spikes from different neurons into a single cluster. We chose this conservative approach deliberately, accepting a higher number of fragments that could later be reunited through manual curation, rather than allowing automatic merging that might produce contaminated clusters. Kilosort outputs were post-processed using SpikeInterface (version 0.102) and a custom manual curation GUI. Because the conservative sorting parameters frequently split single neurons into multiple fragments, particularly when spike amplitude drifted across the recording, oversplit units were identified and merged back together based on template similarity, spatial proximity, cross-correlogram refractory structure, and amplitude continuity. Every unit was individually reviewed in the GUI, which displayed amplitude drift across the full recording (split by pre-sleep, sleep, and post-sleep epochs), mean waveforms from early, middle, and late time segments, cross-correlograms, spatial positions on the probe, and amplitude distributions. Units were accepted, rejected, split, or merged with nearby units. Across the four recordings used for drift analysis, manual curation yielded 109, 130, 204, and 223 accepted units. Of these, 34–52% consisted of multiple merged Kilosort templates (mean 2.2 templates per unit, range 1–11), reflecting the degree of oversplitting by the conservative sorting parameters. Median ISI violation rate (< 1.5 ms refractory period) was below 0.03% across all recordings, with the 95th percentile remaining below 6%, confirming good single-unit isolation. Median firing rates ranged from 0.3 to 2.9 Hz across recordings. For the drift decoder, to guard against confounding drift estimates with unit loss/gain, an additional stability filter removed units with mean firing rate below 0.05 Hz in either the first or second half of the recording, reducing NP_231018_day1 from 130 to 100 units; the other recordings were unaffected.

### Probe trajectory identification and channel assignment to anatomical tissue locations

To assign brain region labels to each recording channel, we used lipophilic dye-assisted probe localisation: After transcardial perfusion-fixation with phosphate-buffered saline (0.1 M) and 4% paraformaldehyde (PFA), brains were extracted and stored in 4% PFA at 4°C overnight. Brains were then embedded in 4% agar, and imaged using Brainsaw: serial 2P-tomography (Ragan et al., 2012) using ScanImage (Vidrio Technologies) and BakingTray (Campbell, 2020). Images were stitched using StitchIt (Campbell et al., 2020) and then registered to the Kim Atlas (25 um resolution) using brainreg (Niedworok et al., 2016; Tyson et al., 2022), BrainGlobe Atlas API (Niedworok et al., 2016). Probe tracks labelled with DiI, DiO and/or DiD were traced using brainglobe-segmentation (Tyson et al., 2022). The probe tracks were aligned to a common reference framework using brainrender (Claudi et al., 2021).

While the angle of anatomical tracts can be precisely determined by tract tracing, to further improve our precision of probe depth, and assign anatomical labels to shanks where anatomical identification was less confident, we computed power spectral density from artifact-free LFP segments (Welch’s method, 2 s Hann windows, 50% overlap, 500 Hz sampling rate) and extracted eight features per channel: relative power in seven frequency bands chosen to avoid 50 Hz mains harmonics with 5 Hz margins (delta 1–4, theta 4–8, alpha 8–13, beta 13–20, low gamma 30–45, high gamma 55–95, HFO 105–145 Hz) plus normalized depth within each shank. An XGBoost binary classifier (max depth 5, learning rate 0.1, class-weight-balanced) was trained to discriminate piriform cortex from non-piriform regions (DEn, Cl, AIV, AID, LO) using histological ground truth from 9 recordings (2,586 channels). Per-shank Viterbi decoding with a fixed transition penalty enforced spatial contiguity of predictions along the probe axis. Leave-one-recording-out cross-validation yielded 79.8% strict accuracy and 90.5% accuracy with ±100 µm spatial tolerance. For recordings with histological probe track reconstructions, sublayer boundaries were refined by warping brainsaw atlas annotations to match LFP-predicted piriform/non-piriform transition depths, preserving the histologically determined sublayer order and relative thickness while correcting for depth registration errors. Short anatomically implausible regions (< 30 µm) were absorbed into neighbours. Channels without histology retained the binary LFP prediction. Each unit was assigned a piriform probability based on its peak channel’s classifier score; units with piriform > 0.5 were classified as piriform cortex.

### Time-varying drift decoder

To track how the population readout changes over a recording session, we trained a time-varying linear decoder whose weight matrix evolves smoothly over the session:

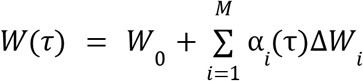

Where *W*_0_ is the baseline weight matrix, 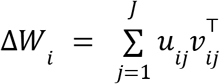 is a low-rank (rank J=3) weight perturbation matrix for mode *i, and* α_*i*_ is a normalised sigmoid activation. The session time τ ∈ [0, 1] maps linearly from the first to the last odour presentation. Each sigmoid activation is normalised to ensure α_*i*_ (0) = 0 and α_*i*_ (1) = 1:

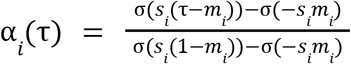

Where *m*_*i*_ is a learnable midpoint (constrained to [0, 1]) and *s*_*i*_ is the slope. This parametrisation means each mode transitions from inactive to fully active once during the session, at a time and rate determined by the data. The decoder input at each 100 ms time bin is the concatenated spike count vector across all N recorded units and C=9 context bins with L=-3 bins of lag (900 ms window, spanning −300 to +600 ms relative to the current bin):

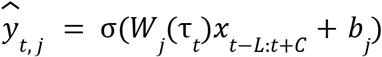

where σ is the logistic sigmoid, *W*_*j*_ is the j-th column of W, and *b*_*j*_ is a learnable per-odour bias.

Each odour is decoded independently (using binary cross-entropy), rather than via a mutually exclusive softmax, because multiple odours can be presented in rapid sequence and the decoder must handle overlapping temporal responses. Based on preliminary experiments with 10 modes (where group lasso regularisation and periodic hard pruning consistently eliminated all but 4–5 modes), the final models were trained with M = 4 modes directly.

### Drift decoder training

Models were trained on all wake periods (NREM bins excluded from training) using AdamW (learning rate 0.001, weight decay 0.001) with cosine annealing over 500 epochs and early stopping (patience 80 epochs on validation loss). Every 6th wake odour presentation per odour was held out for validation. Units were required to have a minimum firing rate of 0.05 Hz in both the first and second halves of the recording. The loss function combines binary cross-entropy (with class-imbalance weighting) and four regularisation terms: group lasso on mode amplitudes (λ_amp_= 0.0046), pairwise redundancy penalty between modes (λ_redund_ = 0.036), within-mode orthogonality (λ_ortho_ = 0.00023). While BCE with positive class weighting optimises odour-detection sensitivity, it tolerates a non-zero prediction floor; we included a background floor penalty (λfloor = 2.4) to suppress tonic predictions during inter-trial intervals. The floor penalty was applied to the lower 68th percentile of non-stimulus predictions, so that rare high-probability events during NREM sleep (potential replay) are not penalised. Midpoints for the first and last modes were frozen during a 60-epoch burn-in to prevent temporal collapse, then freed.

### Drift mode decomposition

After training, each mode’s full weight perturbation was reconstructed as 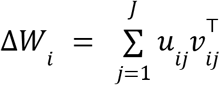 from the learned U and V parameters. These matrices have the same dimensionality as *W*_0_ (N times C rows by 8 odour columns) and can be reshaped to (units, context bins, odours) for visualisation. The gain component of each mode measures the cosine similarity between the vectorised mode weight change and the baseline weights: *gain*_*i*_ = *cos*(*vec*(Δ*W*_*i*_), *vec*(*W* _0_)). A value of 1 would indicate pure scaling of the existing readout; 0 indicates a purely orthogonal rotational change. For reference, the expected absolute cosine between random vectors in d-dimensional space is 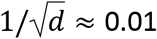 for our weight matrices. The rotation angle per mode was computed as the angle between the baseline and mode-perturbed weight matrices: θ_*i*_ = *arccos*(*cos*(*W*_0_ + Δ*W*_*i*_)), expressed in degrees.

The decorrelation effect of each mode was quantified as the change in mean pairwise Pearson correlation between odour weight vectors when adding the mode: 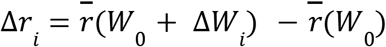 where 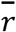 is the mean of the upper triangle of the 8×8 odour correlation matrix. Negative Δ*r* indicates decorrelation (increased odour distinctness); positive indicates dedifferentiation.

The cumulative trajectory of *W*(*τ*) through weight space was visualised by projecting into a 2D subspace via SVD. The four mode weight matrices Δ*W*_1_,…, Δ*W*_4_ (each flattened to a vector) were stacked into a 4 x *d* matrix and decomposed via truncated SVD to obtain the top-2 right singular vectors spanning the plane that best captures variance among the mode directions. No centering was applied, preserving the geometry of angles and distances relative to the origin (*W*_0_). The full trajectory was then computed by evaluating ∑_*i*_α_*i*_(τ) *proj*(Δ*W*_*i*_) at each τ using the actual sigmoid activations, producing a continuous curve from *W*_0_ to *W*_*end*_ that passes through intermediate states at the rate determined by the learned mode dynamics. Each recording was projected independently because the SVD basis is not comparable across recordings with different numbers of units. The 2D trajectory was rotated so that the total drift direction (*W*_0_ → *W*_*end*_) points horizontally, and reflected so that the majority of the path curves upward.

### TDLM sequenceness analysis

To test whether replay events followed the temporal order of odour sequences experienced during wakefulness, we applied Temporally Delayed Linear Modelling (TDLM; Liu et al., 2021). For each sequence (F1 and N1), we constructed a transition matrix *T* by ridge regression (α = 1. 0) predicting the decoder output at lag 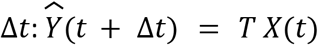, where *X*(*t*) and 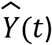 are the 4-odour decoder probability vectors for the relevant sequence at time *t*. The forward sequenceness score was computed as the mean of forward transitions *T* _*i*→*i*+1_ minus backward transitions *T*_*i*+1→*i*_ along the trained sequence order. During wake, the inter-odour onset interval was 1 second (10 bins at 100 ms). We tested replay at compression factors of 1×, 2×, 3×, and 4×, corresponding to lags of 10, 5, 3, and 2 bins, respectively. Significance was assessed against a circular-shift null distribution (500 shuffles). For each shuffle, the decoder prediction time series for each odour channel was independently circular-shifted by a random offset within the NREM mask, preserving the autocorrelation structure and marginal statistics of each channel while destroying inter-channel temporal relationships. The sequenceness score from the real data was compared to the shuffle distribution to obtain a p-value. Bootstrap confidence intervals (95%) were computed by resampling NREM time bins with replacement (500 resamples), recomputing the transition matrix and sequenceness score each time. NREM and wake sequence lengths were compared using matched sample sizes: the number of valid (non-excluded) bins was equalised between states before computing transition matrices, thereby preventing statistical artefacts from unequal sample sizes.

### Field potential analysis

Local field potentials were obtained by low-pass filtering and downsampling the Neuropixels wideband signal to 500 Hz. Artifact detection was run on the raw signal to identify saturated segments (standard deviation < 2.0 µV in 20 ms sliding windows on any channel), large transients (> 3000 µV), step artifacts (> 500 µV per sample), and flat segments (< 0.5 µV). A boundary of 300 ms was expanded around each artifact identified. For all subsequent analyses, only artifact-free segments were used.

Sharp waves (SPWs) were detected in the 5–20 Hz bandpass-filtered LFP (3rd-order Butterworth) using a dual-threshold approach. The envelope of the filtered signal was z-scored against the distribution of clean NREM segments only, so that detection thresholds reflect NREM-specific amplitude statistics rather than being diluted by lower-amplitude wake activity. Events were detected when the z-scored envelope exceeded 1.5 SD (edge threshold), and each event required that the peak within it exceed 3.0 SD. Events shorter than 30 ms or longer than 500 ms were discarded. Events within 20 ms of each other were merged, and a 100 ms refractory period was enforced. Both negative and positive polarity deflections were detected separately. Spatiotemporal coincidence across channels was required: events detected on fewer than 3 channels within the merge window were discarded, ensuring that detected SPWs reflect population-level field potential events rather than single-channel noise. A post-hoc filter selected high-confidence NREM sharp waves: negative polarity, global z-score between 5 and 10 (excluding extreme outliers), duration ≤ 200 ms, and local z-score < −3.0 computed in a ±20 s sliding window. The local z-score requirement ensures that detected events are large relative to their immediate temporal context, not just relative to the overall NREM distribution.

For each detected sharp wave, the presence of co-occurring fast oscillations was assessed in three additional frequency bands: slow gamma (30–40 Hz), fast gamma (80–140 Hz), and ripple (160–190 Hz). Hilbert envelopes for each band were computed and normalised to a ±300 ms baseline (the first and last 100 ms of the peri-event window). Peri-SPW-triggered averages were computed across all NREM events passing the post-hoc filter, with 95% confidence intervals (mean ± 1.96 SEM).

To test whether replay events preferentially co-occurred with sharp waves, we computed the probability of observing a replay event within 0–500 ms following each sharp wave. Replay events were identified as time bins during NREM sleep where the drift decoder produced above-threshold odour predictions in the absence of odour stimulation. At each time bin *t*, the decoder output 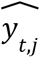 gives the probability that odour *j* is present, using the time-appropriate weights *W* _τ_. A replay event was defined as any NREM bin where 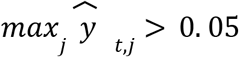. This threshold was chosen to be well above the background prediction floor (median < 0.01 during NREM) while remaining sensitive to weak reactivation. To avoid contamination from stimulus-evoked responses, bins within a 5-second halo of any odour presentation were excluded from both NREM and wake replay analyses.

## Acknowledgements

We thank Kayvan Combadiere for help with setting up the spikesorting pipeline; Dan Gunton for help with implementing Brainsaw; George Konstantinou for help with designing custom PCBs; the Crick Mechanical Engineering Team for help with manufacturing custom bottle-holders and manifolds; and Goncalo Lopes for help with our Bonsai workflow. We also thank Carl Schoonover and Andrew Fink for helpful discussions about drift and our data; Kris Jensen and Tim Behrens for helpful advice regarding replay analysis; and Cecilia Della Casa and Sina Tootonian for valuable help regarding the drift modelling approach.

---

**Supplementary Figure 2.**
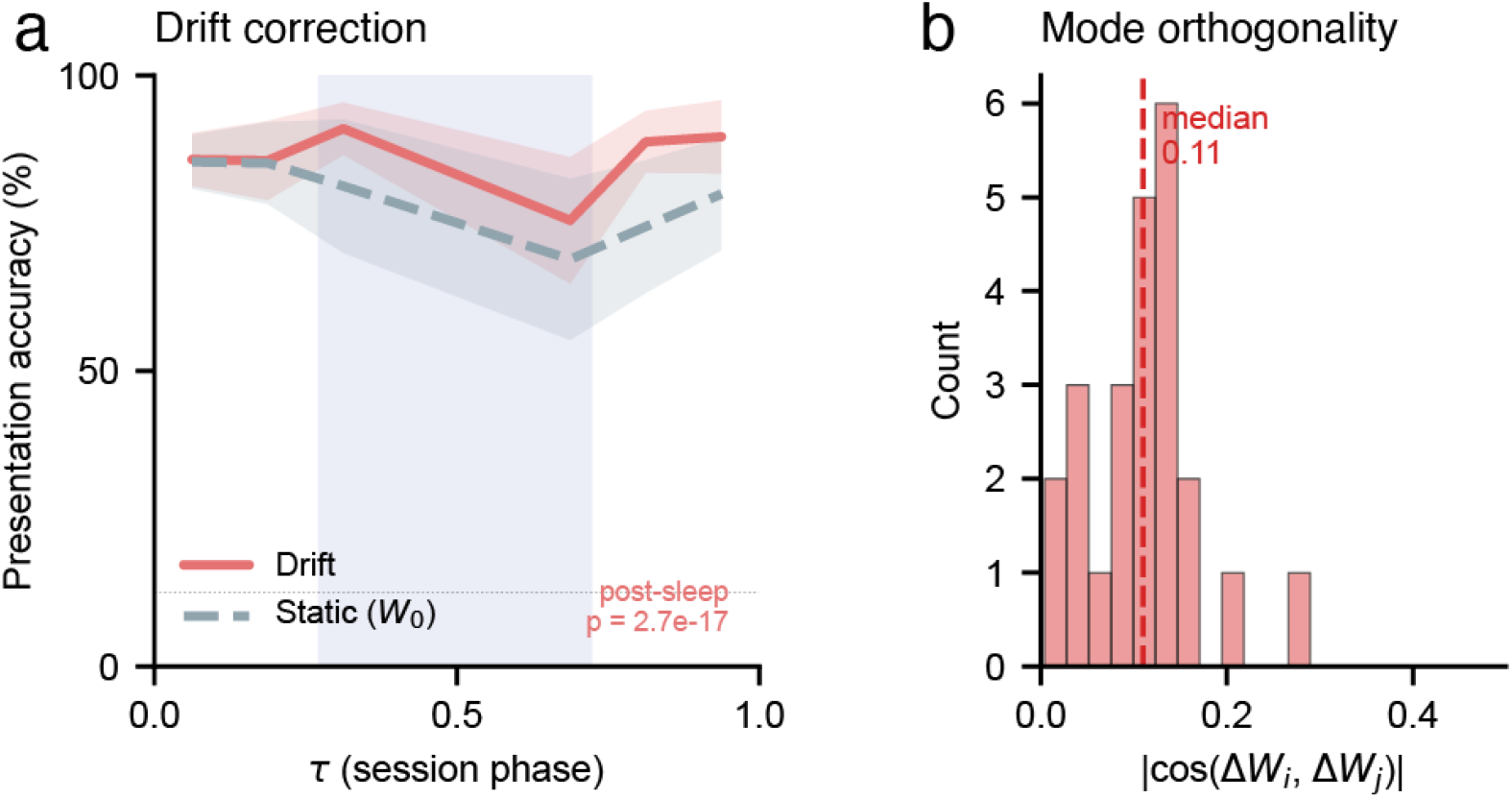
Drift mode impact and orthogonality. **(a)** Per-presentation decoding accuracy across the session for the drift-corrected decoder (red) versus a static decoder using frozen baseline weights *W*_0_ (grey dashed). Accuracy is computed per odour presentation (1 s average of decoded probabilities, argmax classification) and binned by session phase τ (8 bins). Lines and shading show mean ± SEM across 4 mice. Both decoders perform comparably during pre-sleep wake, but the static decoder degrades post-sleep as the population representation drifts away from *W*_0_, while the drift decoder maintains accuracy by tracking the evolving readout. Dotted line: chance level (12.5%, 1/8 odours). Blue shading: mean NREM sleep period. Post-sleep improvement is significant (pooled McNemar test across all post-sleep presentations: 119 presentations correctly classified only by the drift decoder vs 22 only by the static decoder, p = 2.7 × 10^−17^). **(b)** Pairwise orthogonality of the four drift modes. Histogram of absolute cosine similarity for all 6 mode pairs across 4 recordings (24 values total). For reference, perfectly orthogonal vectors would give |cos| = 0; random vectors in the same dimensionality would give expected |cos| ≈ 0.01. The modes are nearly orthogonal, confirming that each captures an independent component of representational drift.

**Supplementary Figure 3.**
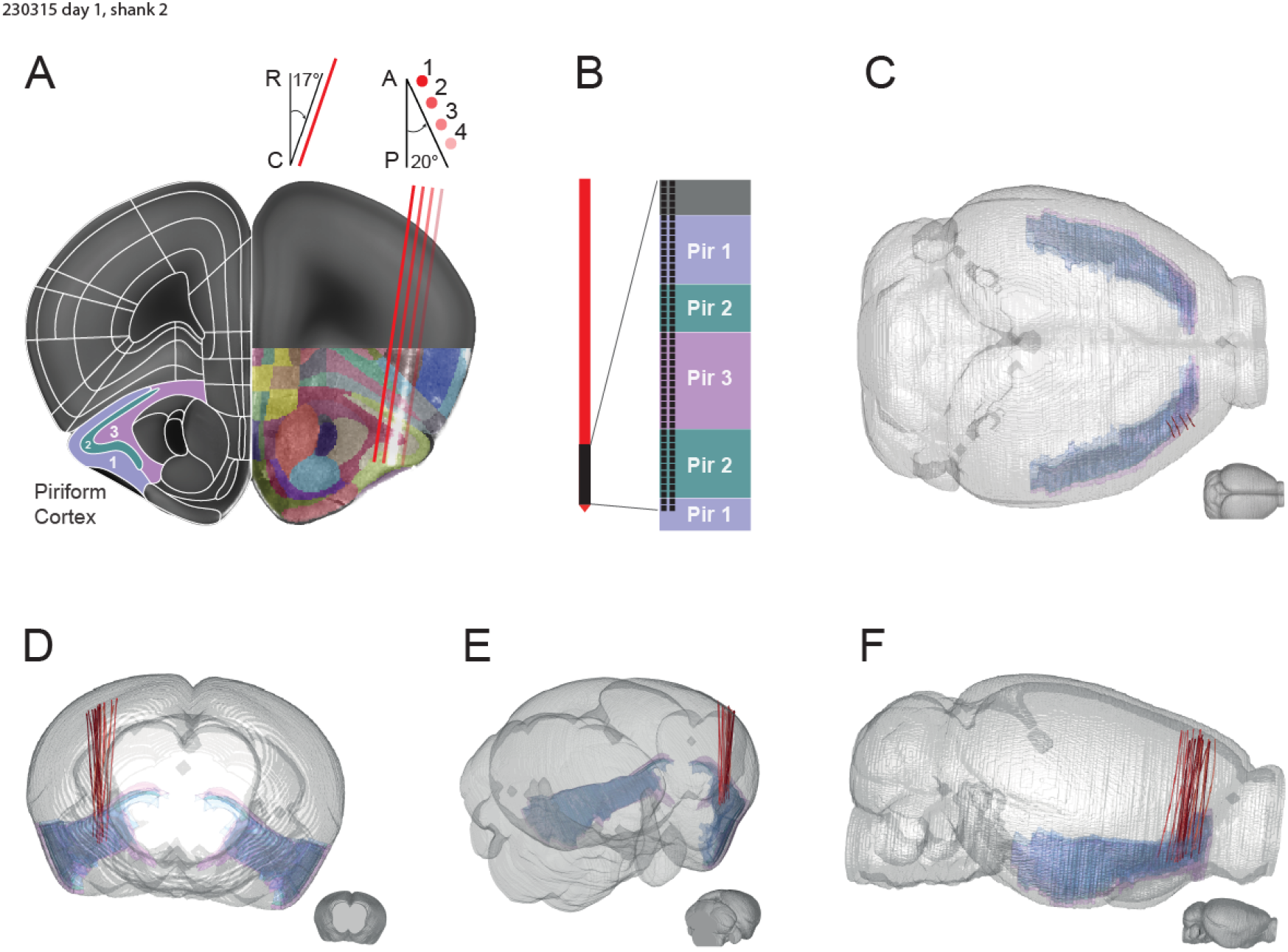
Probe localisation. **(a)** Coronal view of Kim Atlas (left hemisphere) and multi-shank probe track (right). Other probes are in front or behind the plane of focus due to insertion angle (top). **(b)** Active channels and their position in the brain for an example shank. **(c)** Horizontal view showing how probe angle follows the shape of the piriform cortex. **(d-f)** Probe locations of all insertions.

**Supplementary Figure 4.**
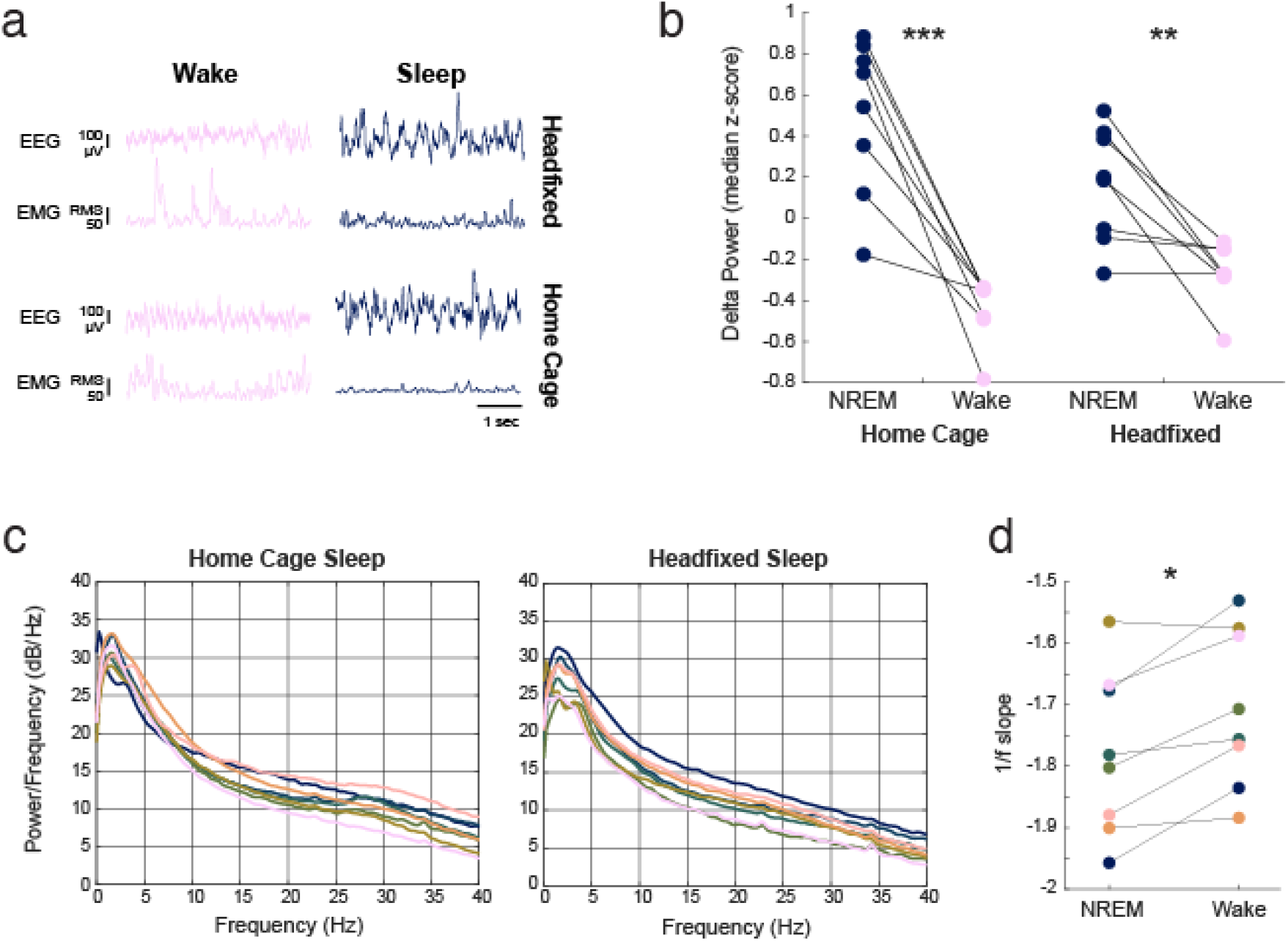
Headfixed sleep. **(a)** Example raw EEG and EMG traces from headfixed wake and sleep (top row) in comparison to home cage wake and sleep (bottom row) for the same mouse. **(b)** Delta power during wake and sleep in home cage vs headfixed for all mice. **(c)** Power spectra during sleep in home cage (left) is comparable to headfixed sleep (right). **(d)** 1/f slope is steeper during sleep than wake (all headfixed).

